# A Hybrid Type I and II Polyketide Synthases Yields Distinct Aromatic Polyketides

**DOI:** 10.1101/2024.08.28.610196

**Authors:** Li Ya Zhao, Jing Shi, Zhao Yang Xu, Jia Lin Sun, Zhang Yuan Yan, Zhi Wu Tong, Ren Xiang Tan, Rui Hua Jiao, Hui Ming Ge

**Author notes:** **Corresponding Author** (J.S.);, (R.H.J.);, (H.M.G.). **Author Contributions** L.Y.Z. and J.S. contributed equally to this work.

## Abstract

Bacterial aromatic polyketides are compounds with multiple aromatic rings synthesized by bacterial type II polyketide synthases (PKSs), some of which have been developed into clinical drugs. Compounds containing aromatic polyketides synthesized by a hybrid type I and type II PKSs are extremely rare. Here, we report the discovery of a gene cluster encoding both modular type I PKS, type II PKS and KAS III through extensive bioinformatics analysis, leading to the characterization of the hybrid polyketide, spirocycline A. The structure of spirocycline A is unprecedented among all aromatic polyketides, featuring a unique starter unit, four spirocycles, and forming a dimer. Biosynthetic studies indicate that the starter unit of this molecule is synthesized by type I PKS in collaboration with two *trans*-acting ketoreductase (KR) and enoylreductase (ER). It is then transferred by KAS III to the type II PKS system, which then synthesizes the tricyclic aromatic polyketide backbone. The subsequent formation of the spirocycle and dimerization is carried out by four redox enzymes encoded in the gene cluster. Overall, the discovery of spirocycline A provides a new approach for identifying novel aromatic polyketides and offers potential enzymatic tools for the bioengineering of these hybrid polyketides.

**Table of Contents:** 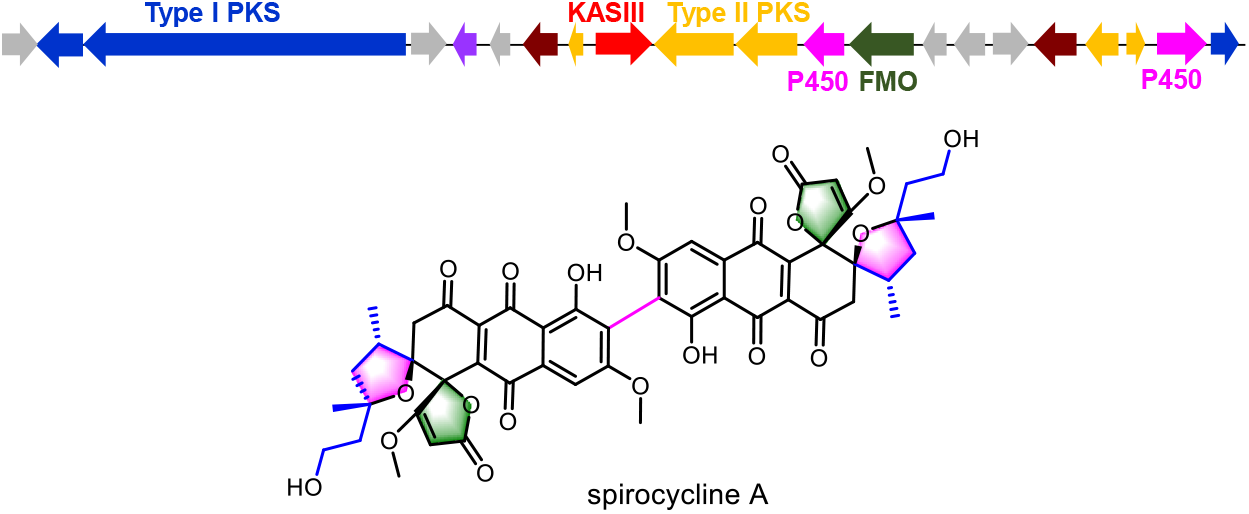

## INTRODUCTION

Bacterial aromatic polyketides, characterized by polycyclic aromatic ring systems, represent a significant category of natural products with distinctive structures.^1-2^ Many of these polyketides have exhibited excellent bioactivities and have been developed into clinical drugs, such as the well-known antibiotic tetracycline and the anticancer agents daunorubicin.^3-4^ These compounds are synthesized by type II polyketide synthases (type II PKSs), which catalyze iterative Claisen condensation reactions to generate a polyketide chain.^5^ This chain then undergoes cyclization and aromatization under the action of ketoreductases (KRs), cyclases (CYCs), and aromatases to form an aromatic ring system. Subsequent modifications by tailoring enzymes such as oxidoreductases, methyltransferases, and glycosyltransferases lead to the structural diversity observed in this class of natural products.^1^

Typically, most bacterial aromatic polyketides utilize acetyl-CoA as the starter unit, but a minority also employ non-acetate starter units such as propionyl-CoA, butyryl-CoA, or even short fatty acyl as starter unit,^6^ as seen in compounds like R1128,^7^ benastatin,^8^ alnumycin^9^ and aclacinomycin^10-11^ (Figure 1A). The incorporation of these non-acetate start units requires the mediation of an additional 3-ketoacyl-ACP synthase (KAS III).^12^ KAS III was initially discovered in fatty acid biosynthesis and is responsible for forming a C-C bond between acetyl-CoA and malonyl-ACP in the first step of fatty acid biosynthesis.^13^ Homologous enzymes of KAS III in type II PKS pathway play a key role as a gatekeeper in selecting specific non-acetate starter unit into the assembly line,^14^ thus greatly increasing the structural diversity and complexity of bacterial aromatic polyketides.

**Figure 1.**
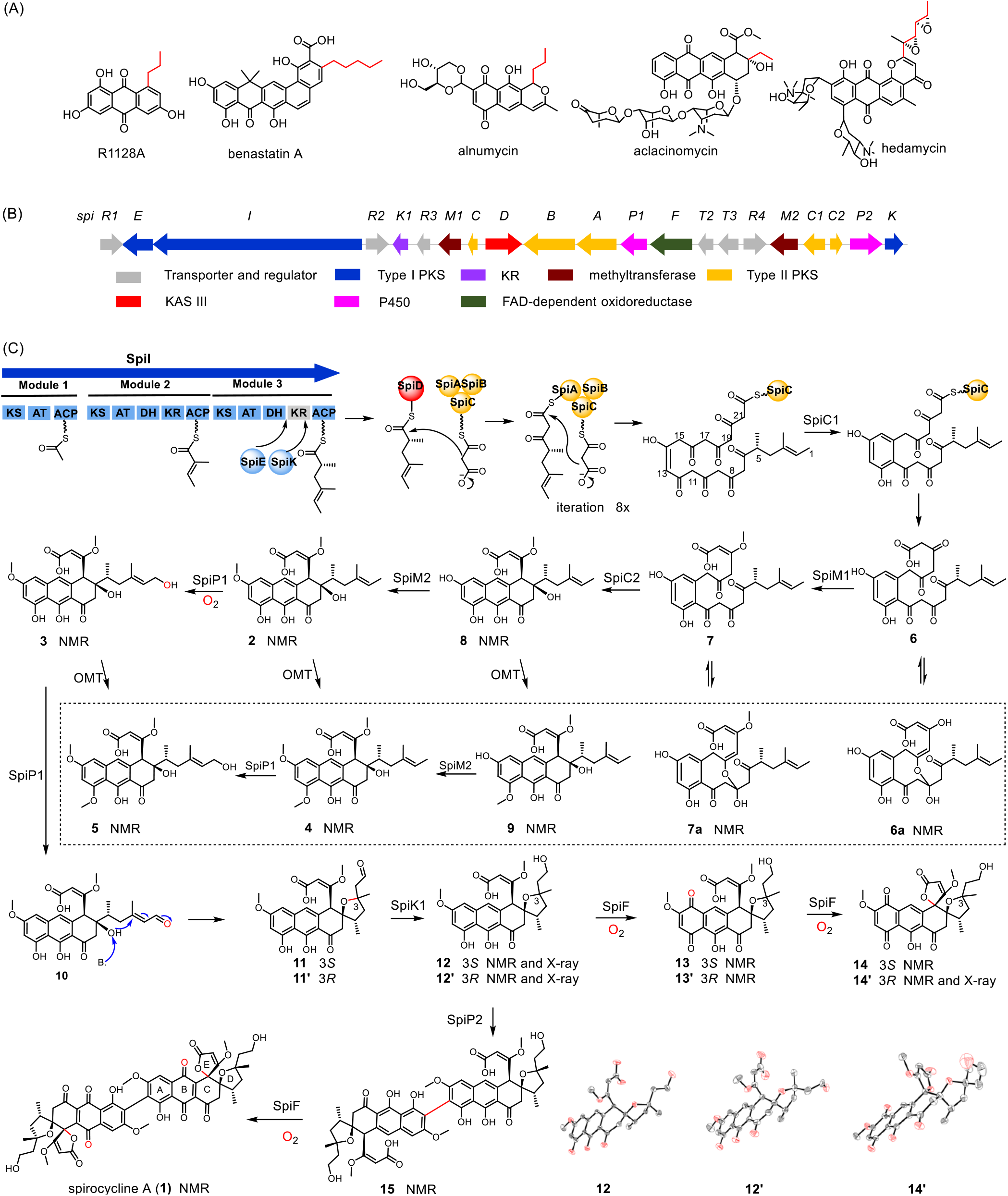
(A) Examples of natural products synthesized by type II PKSs utilize unusual starter units (highlight in red). (B) Biosynthetic gene cluster of *spi* from *S. spectabilis* NA07643. (C) Proposed biosynthetic pathway for spirocycline A (**1**). ORTEP diagram for the crystal structures of **12, 12’** and **14’** are shown in the figure, with CCDC numbers being 2348611, 2348612, and 2348613, respectively.

Among aromatic polyketides that utilize unusual starter units, hedamycin (Figure 1A), a highly selective DNA-alkylating agent, stand out as a unique example.^15^ The starter unit for hedamycin is a hexenoyl acyl unit synthesized by an iterative type I PKS. This unusual starter polyketide then primes the type II PKS through the mediation of a KAS III. To date, only one example of such hybrid type I and II polyketide has been reported. Compared to fatty acyl groups, the functional groups derived from type I PKS can be further modified by post-modification enzymes. For example, the two double bonds on the starter unit of hedamycin can be further oxidized to form two epoxide rings. Notably, the introduction of polyketide starter units not only increases the structural complexity but also significantly increase the antitumor activity of hedamycin.^16^

Herein, we conducted a systematic analysis of type II PKS clusters in public genome databases and identified a class of gene clusters encoding both type I and type II PKSs, as well as KAS III, with multiple genes encoding post-modification enzymes. By optimizing fermentation and cultivation conditions, we obtained a novel aromatic polyketide with unprecedented structural features. Through *in vivo* and *in vitro* studies, we elucidated its biosynthetic process, which begins with the biosynthesis of an unusual 2,4-dimethylhex-4-enoic acyl starting unit synthesized by a modular type I PKS, followed by its transfer to type II PKS mediated by KAS III. Under the action of cyclases and methyltransferases, an aromatic polyketide monomer is formed, which is then further processed by multiple oxidoreductases to form two unique spiro rings and dimerize into the final product spirocycline A (**1**).

## RESULSTS AND DISCUSSION

### Identification of the hybrid PKS gene cluster

Previous genome mining of type II PKS has primarily focused on the analysis of CLF and post-modification oxidoreductase.^17-19^ While this approach continues to yield new aromatic polyketides with varying chain lengths and unusual oxidative modifications, the discovery potential is restricted to known carbon skeletons.^19-20^ To uncover novel molecules, we aim to mine type II PKS with unique starter molecules. To this end, we analyzed a total of 21,728 annotated genome files of actinomycetes from the NCBI database as well as our own sequenced data. Among them, there were 5,944 type II PKS gene clusters. After further screening, we obtained 919 type II PKS clusters containing KAS III. These gene clusters were systematically analyzed, and two homologous gene clusters (*mic* and *spi*) from *Micromonospora* sp. HM134 ^21^and *Streptomyces spectabilis* NA07643, respectively, were selected for further characterization based on the co-occurrence of type I and type II PKSs and KAS III genes (Figure S1). It is speculated that these two clusters are likely responsible for synthesizing hybrid polyketides, with their starting units most likely being Type I polyketides rather than the commonly found fatty acids.

We fermented *M*. sp. HM134 and *S. spectabilis* NA07643 strains on different media (Figure S2). Through LC-MS analysis of the fermentation products, we found that the *S. spectabilis* NA07643 strain could produce five products, **1**-**5** (Figure 2A i), in the RM fermentation medium, while its corresponding KS mutant strain Δ*spiA* was not able to afford these products (Figure 2A ii), confirming that they are related to the *spi* gene cluster. However, *M*. sp. HM134 strain did not produce any similar UV-absorbing compounds during the screening on various media (Figure S2). Therefore, we carried out a 20-L scale fermentation of *S. spectabilis* NA07643 on RM medium and successfully obtained **1**-**5**.

**Figure 2.**
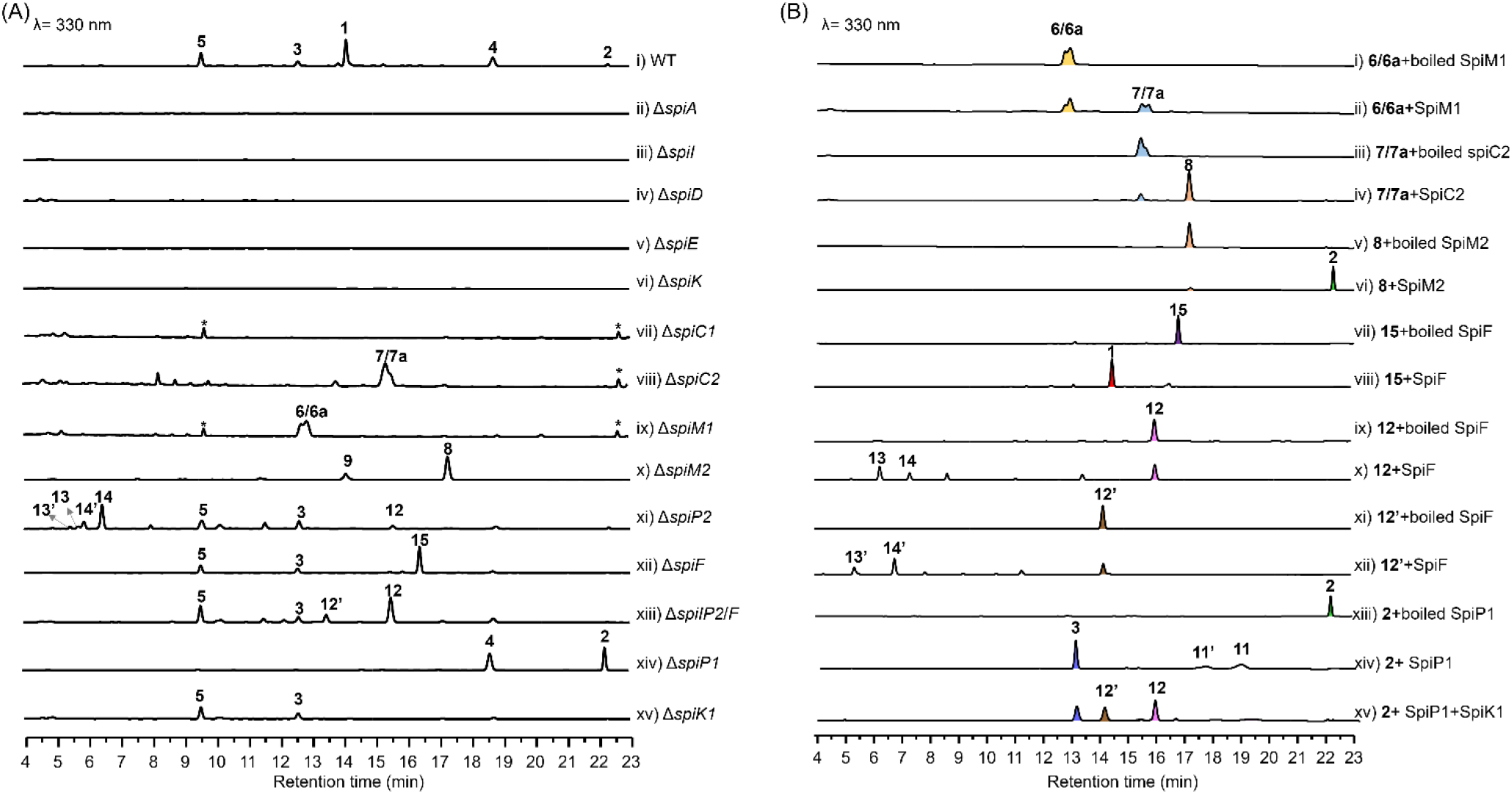
(A) HPLC analysis of metabolites from wild-type (WT) strains and mutant. (B) HPLC analysis of organic extracts from different reconstituted *spi* enzyme sets.

Compound **1** was isolated as an orange powder. The molecular formula was determined to be C_52_H_50_O_20_ based on its HRESIMS. However, its ^13^C NMR spectrum only showed 26 carbon signals, suggesting that **1** is a symmetric dimer. Based on the 1D and 2D NMR analysis, **1** is structurally determined as an anthraquinone-type polyketide dimer with two pairs of unique spiro-rings (Table S5). The relative and absolute configurations of **1** were later established by comparing them with those of **12, 12’**, and **14’** (Figure S3), which were isolated from the mutant strains and determined by X-ray diffraction (Figure 1C). Compounds **2**-**5** were structurally determined as monomeric compounds that have not yet formed spirocycles (Tables S6-S9). When **2**-**5** were fed to the Δ*spiA* mutant strain, only **2** and **3** were able to restore the production of **1**, confirming that they are on-pathway biosynthetic intermediates (Figure S4). In contrast, **4** and **5** were identified as shunt products.

### Biosynthesis of the monomeric intermediate 2

Based on the structure of **2**, it can be speculated that its biosynthesis starts with 2,4-dimethylhex-4-enoic acyl triketide, followed by eight rounds of extension under the action of type II PKS. Subsequently, CYCs and methyltransferases (MTs) are involved in the cyclization and methylation processes to form **2**. In the *spi* gene cluster, *spiI* encodes a type I PKS with three modules, which is likely responsible for catalyzing the formation of the triketide starting unit. Bioinformatic analysis shows that each domain in SpiI is functional, except for the KR domain in module 3, which lacks the critical catalytic residues K and Y(Figure S5).^22^ Additionally, module 3 is missing an enoylreductase (ER) domain. However, the *spi* gene cluster encodes a standalone KR (SpiK) and an ER protein (SpiE), which share 34% and 37% sequence identity with the FabG (KR) and FabV (ER) domains of fatty acid synthase,^23-24^ respectively, suggesting that these two proteins likely interact with SpiI and compensate for the missing KR and ER functions.

When the *spiI, spiK, spiE, spiD* and *spiC1* genes were individually knocked out, the production of **1**-**5** abolished (Figure 2A iii-vii), indicating that these genes are essential and likely act on the polyketide intermediate before its release from the acyl carrier protein (ACP). When we knocked out *spiM1* (MT), a pair of inseparable products **6** and **6a** with the same molecular weight were generated (Figure 2A ix). Structure elucidation of **6**/**6a** indicated they are a pair of tautomeric isomers in which only the first benzene ring of the tricyclic polyketide framework has been formed (Tables S10-11). Similarly, the Δ*spiC2* (CYC) mutant led to produce another pair of isomers **7** and **7a** (Figure 2A viii), both of which has 14 Da greater than **6/6a**. NMR analysis confirmed **7**/**7a** are the OH-20 methylated derivatives of **6**/**6a** (Tables S12-13), implying that SpiM1 might act before SpiC2 and catalyzes the methylation on enol-20 group. When we further inactivated the second MT encoding gene, *spiM2*, two clear peaks **8** and **9** were generated from the resulting mutant strain (Figure 2A x). Compound **8** lacks the methyl group at OH-14 compared to **2**, indicating SpiM2 is a MT acting on OH-14, while **9** is an off-pathway shunt product with an additional methyl group at 12-OH (Tables S14-15).

To further verify the biosynthetic pathway, we heterologously expressed and purified SpiM1, SpiM2 and SpiC2 in *E. coli* BL21(DE3), respectively (Figure S7). When **6/6a** were incubated together with SpiM1 in the presence of the SAM cofactor, the formation of **7/7a** were detected (Figure 2B i-ii). Upon further addition of SpiC2, **7/7a** gradually disappeared, and a new product **8** was therefore generated (Figure 2B iii-iv). In addition, under the action of SpiM2, **8** can be converted to the sole product **2**, instead of **9**, verifying the function of SpiM2 (Figure 2B v-vi). Thus, we concluded that SpiI, along with SpiK and SpiE, synthesizes the unusual triketide, which initiates type II PKS biosynthesis via SpiD (KAS III). After eight rounds of elongation, SpiC1 catalyzes the formation of the first aromatic ring, resulting in the proposed intermediate **6**. SpiM1 then methylates the enol to form **7**, and SpiC2 cyclizes the second and third rings, creating the tricyclic aromatic polyketide backbone **8**, which is further methylated to produce **2** (Figure 1C).

### SpiP2 catalyzes the oxidative dimerization

We reasoned that the subsequent steps from **2** to **1**, including dimerization, spirocyclization, and oxidation of the naphthalene ring, are likely mediated by the participation of redox enzymes including SpiF (FMO), SpiP1 (P450), and SpiP2 (P450) that are encoded in the gene cluster (Figure 1B). Cytochrome P450 enzymes play a key role in the crosslinking of aromatic rings in various natural products, such as vancomycin,^25^ himastatin,^26^ citillin,^27^ and some recently found P450-modified RiPPs.^28^ Notably, P450 enzymes have also been reported to catalyze the dimerization of aromatic polyketide such as the bacterial aromatic polyketide julichrome^29-30^ and the fungal aromatic polyketide orlandin^31^. We constructed a phylogenetic tree including the P450 enzymes from the *spi* gene cluster and other characterized bacterial P450s (Figure S8). The analysis revealed that SpiP2 and JuiI, the P450 responsible for dimerization in julichrome biosynthesis,^29-30^ are grouped in the same clade, while SpiP1 clusters with several P450 enzymes with hydroxylation functions.

The Δ*spiP2* mutant strain no longer produced **1**, but accumulated nine compounds, **2, 3, 5, 12, 13, 13’, 14**, and **14’** (Figure 2A xi). All of these products were isolated and structurally characterized by NMR analysis (Tables S16-S21), and **12** and **14**’ were further confirmed by X-ray analysis. It is noteworthy that all these isolated compounds are monomers. Among them, **13** and **13’** as well as **14** and **14’** are pairs of diastereoisomers differing in the configuration at C-3. Despite our numerous efforts, we were unable to obtain soluble P450 protein, which impeded the in vitro validation of its function. However, based on phylogenetic analysis and the monomeric compounds produced from Δ*spiP2* mutant, it can be inferred that the function of P450 is related to dimerization.

### SpiF catalyzes a spirocyclic ring formation

FMOs play crucial roles in shaping structural diversity and complexity in type II polyketide biosynthesis.^32-37^ Phylogenetic analysis revealed that FMOs in type II PKS biosynthesis can be mainly divided into four clades (Figure S9). Clade 1 comprises FMOs that catalyze hydroxylation at the C1 position, such as OxyE in oxytetracycline biosynthesis,^32^ while Clade 2 are FMOs that catalyze hydroxylation at the C4a position, like OxyL^.33^ SpiF belongs to clade 3, whose members along with those in clade 4 display diverse functionality, including catalyzing Baeyer-Villiger oxidation,^34-35^ hydroxylation,^36^ epoxidation, and so on.^37^

To evaluate the role of SpiF in the biosynthesis of **1**, we generated Δ*spiF* mutant, which no longer produced **1**. Instead, a new product **15** was accumulated as a dominant product along with **3** and **5** (Figure 2A xii). Structural analysis revealed that **15** is a dimeric form of **12** (Table S22), indicating that SpiF likely acts after the dimerization process, with compound **12** being the substrate of SpiP2. To verify this hypothesis, we constructed a double knockout strain, Δ*spiF/P2*. In comparison to the Δ*spiP2* mutant strain, **13, 13’, 14** and **14’** were no longer produced. Instead, the yield of **12** was significantly increased as the dominant product (Figure 2A xiii), accompanied by a new compound **12’** with the same molecular weight. Structural elucidation revealed **12’** to be a diastereomer of **12** (Table S17), differing in the configuration at C-3, which was further supported by X-ray crystallographic analysis (Figure 1C).

It is noteworthy that the second 5/6-spirocyclic ring failed to form in all compounds accumulated in Δ*spiF* mutant strain, suggesting that SpiF is likely responsible for catalyzing this spiro-ring formation. To verify the function of SpiF, we overexpressed SpiF protein from *E. coli* BL21(DE3). SpiF is a bright yellow protein, and the LC-MS analysis of the denatured protein supernatant confirmed the presence of FAD cofactor (Figure S10). Upon incubation of **15**, SpiF, and NADPH, we detected the production of **1** (Figure 2B vii-viii). Therefore, SpiF catalyzed not only the oxidation of the naphthyl ring but also the formation of the spirocyclic ring. Furthermore, when SpiF was assayed with **12** under the same reaction conditions (Figure 2B ix-xii). The LC-MS analysis of the reaction mixture showed that SpiF can act on **12**, initially hydroxylating it to produce **13**, which subsequently converts into the spirocyclic compound **14** (Figure 1C). Also, SpiF can catalyze a similar reaction using **12’** to form **14’** via **13’** (Figure 1C).

In addition, during the time course of the enzymatic reaction with SpiF using **15** as the substrate, we observed the ion signal corresponding to the monohydroxylated naphthyl compound **15a**, with a mass increase of 16 Da compared to **15** within 2 minutes. After 4 minutes, the ion signals for the putative dihydroxylated derivative **15b** and monocyclized **15c** were detected (Figure 3B). The formation of **1** was observed at the 6-minute mark. Ultimately, after 8 minutes, all substrates and intermediates were fully consumed and converted into **1**. Based on these results, a mechanism for spirocyclic ring formation is proposed as shown in Figure 3A. SpiF initially oxidizes the naphthyl ring, leading to the formation of a hydroquinone compound either on ring A (monomeric species) or ring B (dimeric species). This hydroquinone then undergoes spontaneous oxidation to the corresponding *o*-quinone. Following tautomerization, the *o*-quinone can transform into **15d** which then engages in an intramolecular Michael addition to forge the spirocyclic scaffold. The resulting hydroquinone is again oxidized to the corresponding *o*-quinone form in **1**.^38^

**Figure 3.**
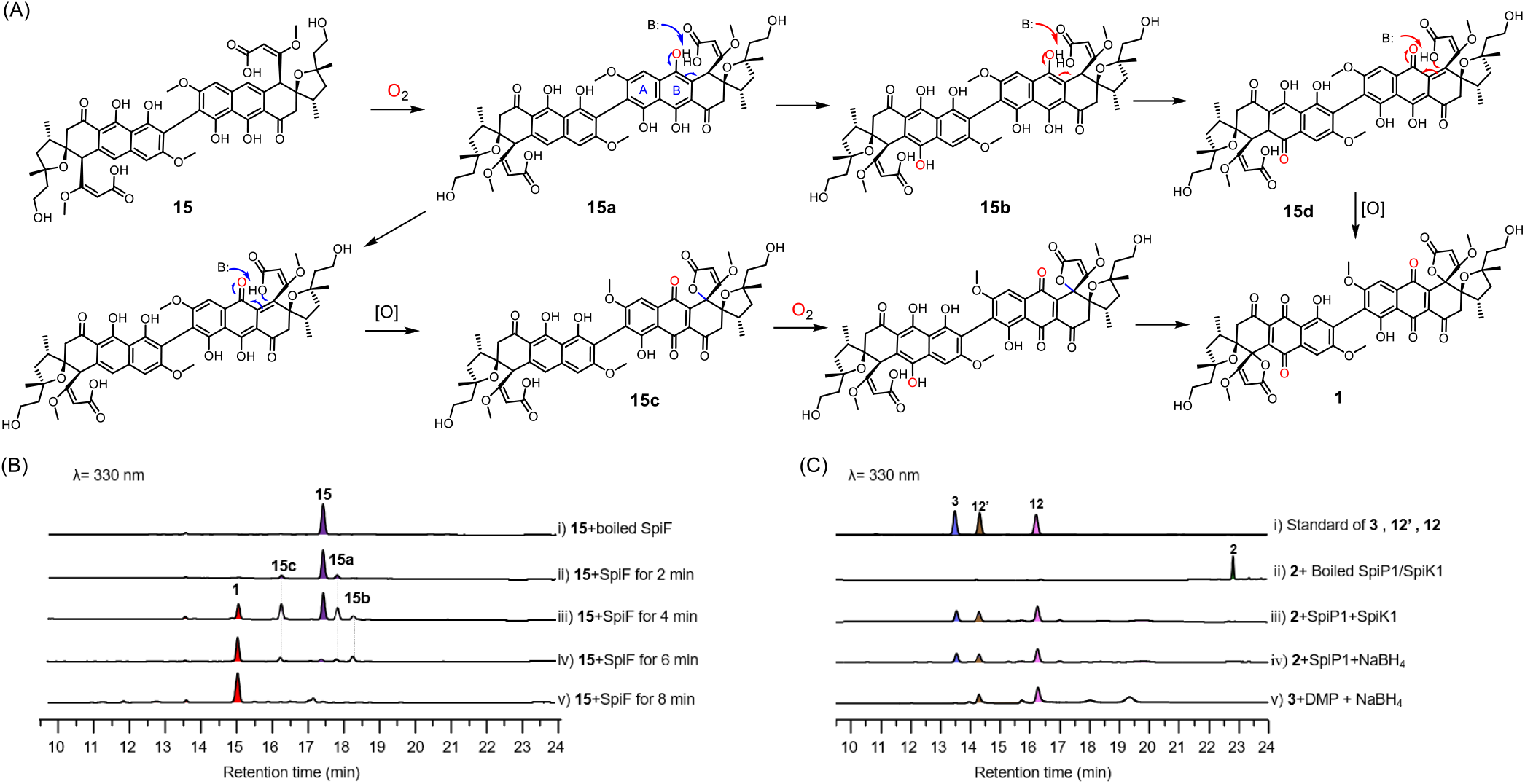
(A) Proposed mechanism for SpiF-catalyzed reaction. (B) In vitro biochemical reactions catalyzed by SpiF. The mass spectrometry data for compounds **15a, 15b**, and **15c** can be found in Figure S139 (C) In vitro biochemical reactions catalyzed by SpiP1/K1.

### SpiP1 and SpiK1 catalyze the second spirocyclic ring formation

When we knocked out another P450 gene, *spiP1*, two compounds, **2** and **4**, were accumulated in large amounts, both featuring a terminal methyl group on their side chains. This suggests that SpiP1 could be involved in the oxidation of the terminal methyl group. We expressed and obtained soluble SpiP1 protein from *E. coli* BL21(DE3) (Figure S7). When **2** was assayed with SpiP1 in the presence of Fdr, Fdx, and NADPH, we could detect the formation of **3** (Figure 2B xiii-xiv). And we also observed the formation of two putative aldehyde products **11**/**11’** with a mass 2 Da less than **3**. However, upon further extending the reaction time, the substrate **2** as well as **3** and **11**/**11’** were gradually consumed, but no new peaks emerged (Figure S11). We reasoned that P450 oxidizes **2** to the aldehyde intermediate **10**, subsequently an intramolecular Michael addition could occur to form the cyclized products **11**/**11’** (Figure 1C). It is conceivable that for the formation of **12**/**12’**, a KR would be required to reduce the unstable aldehyde moiety, thereby yielding the hydroxyl group present in **12**/**12’**.

Within the gene cluster, there is another KR gene, *spiK1*, which shares 37% sequence identity with FabG.^33^ Deletion of *spiK1* led to the production of **3** and **5** (Figure 2A xv), confirming that SpiK1 acts after SpiP1. Incubation of **2** with recombinant SpiK1 and SpiP1 together with the required co-factor and coenzymes led to the successful formation of **12** and **12’** in a ratio of ∼5:2 (Figure 2B xv). In the absence of any one component from the reaction system, **12**/**12’** was no longer produced. Therefore, SpiP1 could oxidize the C-1 position of **2** to give the aldehyde **10**, followed by a spontaneous Michael addition of the C-6 hydroxyl group attacking the C-3 position to form the spirocyclic compounds **11**/**11’**. Subsequently, SpiK1 reduces the aldehyde to a hydroxyl group, yielding **12**/**12’** (Figure 1C). Since aldehyde compounds are toxic to biological systems, in the Δ*spiK1* strain, other KRs in the genome likely reduced the aldehyde group in **10**, thereby yielding **3** and **5**.

To further corroborate the role of SpiK1, we used NaBH_4_ instead of SpiK1 in the above reaction system. Intriguingly, we observed the formation of **12** and **12’** in a ratio consistent with that obtained from the enzymatic reaction (Figure 3C iv). Moreover, when we directly treated **3** with Dess-Martin periodinane (DMP) for 5 minutes, followed by reduction with NaBH_4_, the reaction results showed that **12** and **12’** were successfully generated (Figure 3C v). These results substantiate that SpiP1 solely catalyzes the oxidation of the terminal methyl group to an aldehyde, thereby facilitating an intramolecular Michael addition to forge the spirocyclic scaffold. Subsequently, SpiK1 reduces the aldehyde functionality to the corresponding hydroxyl group (Figure 1C).

## CONCLUSION

In this work, we discovered a gene cluster named *spi*, containing both type I and type II PKS as well as KAS III, through bacterial genome mining. This cluster is responsible for synthesizing an unprecedented aromatic polyketide compound, spirocycline A, which features a unique polyketide starter unit, two pairs of rare spirocyclic rings, and a dimeric form. We elucidated the biosynthetic pathway of this compound, showing that the modular type I PKS, along with KR and ER, synthesizes the special starter unit. This unit is recognized and transferred by KAS III into the type II PKS assembly line, leading to the synthesis of the tricyclic aromatic polyketide **2**. Additionally, we identified four redox enzymes for further structure modificaiton. The P450 enzyme SpiP1 catalyzes the oxidation of the terminal methyl group of **2** to form the aldehyde compound **10**, which triggers a spontaneous Michael addition to create the first spiro ring. The aldehyde group is then reduced by SpiK1 to the hydroxyl compound **12**, which can be utilized by SpiP to give the dimeric compound **15**. SpiF finally oxidizes **15**, completing the construction of the second spiro ring and resulting in a more complex structural scaffold. Given the vast genetic potential of bacterial genomes, more type I and type II PKS hybrid polyketides are likely to be uncovered, holding great promise for synthetic biology and medicinal chemistry. Our study also demonstrates how biosynthetic research can facilitate biocatalytic innovations.

## ASSOCIATED CONTENT

### Supporting Information

The Supporting Information is available free of charge at http://pubs.acs.org.

Experimental procedures and NMR spectra for all isolated compounds (PDF)

### Accession Codes

CCDC 2348611, 2348612, and 2348613 contain the supplementary crystallographic data for this paper. These data can be obtained free of charge via www.ccdc.cam.ac.uk/data_request/cif, by emailing, or by contacting The Cambridge Crystallographic Data Centre, 12 Union Road, Cambridge CB2 1EZ, U.K.; fax:+44 1223 336033.

## AUTHOR INFORMATION

### Author Contributions

† L.Y.Z. and J.S. contributed equally to this work.

### Notes

The authors declare no competing financial interest.

## ACKNOWLEDGMENT

This work was financially supported by MOST (2022YFC2804100 and 2022YFC2303100), and NSFC (22377052, 81925033, 22193071, 81991522, 81991524, and 22107048).

## Notes

### Competing Interest Statement

The authors have declared no competing interest.

